# Confirmation and fine mapping of the resistance locus *Ren9* from the grapevine cultivar ‘Regent’

**DOI:** 10.1101/2020.11.27.400770

**Authors:** Daniel Zendler, Reinhard Töpfer, Eva Zyprian

## Abstract

Grapevine (*Vitis vinifera* ssp. *vinifera*) is a major fruit crop with high economic importance. Due to its susceptibility towards fungal pathogens such as *Erysiphe necator* and *Plasmopara viticola,* the causal agents of powdery and downy mildew (PM, DM), grapevine growers annually face a major challenge in coping with shortfall of yield caused by these diseases. Here we report the confirmation of a genetic resource for grapevine resistance breeding against PM. During the delimitation process of *Ren3* on chromosome 15 from the cultivar ‘Regent’, a second resistance-encoding region on chromosome 15 termed *Ren9* was characterized. It mediates a trailing necrosis associated with the appressoria of *E. necator* and restricts pathogen growth. In this study, we confirm this QTL in a related mapping population of ‘Regent’ x ‘Cabernet Sauvignon’. The data show that this locus is located at the upper arm of chromosome 15 between markers GF15-58 (0.15 Mb) and GF15-53 (4 Mb). The efficiency of the resistance against one of the prominent European PM isolates (EU-B) is demonstrated. Based on fine-mapping and literature knowledge we propose two possible regions of interest and supply genetic markers to follow both regions in marker assisted selection.

## 1. Introduction

The era of accelerated plant breeding started with the emergence of marker-assisted selection (MAS). With this tool in hand, breeders dealing with woody perennials became able to select promising progeny with the desired characteristics at the very early seedling (cotyledon) stage. In grapevine, the requested characteristics are primarily resistance traits against several pathogens, as viticulture worldwide is threatened by a variety of different pests [1,2]. One of the most prominent diseases in vineyards is powdery mildew (PM) caused by the obligate biotrophic ascomycete *Erysiphe necator* (syn. *Uncinula necator* (Schw.) Burr; anamorph *Oidium tuckeri* Berk). This pathogen occurs predominantly in dry and warm regions. *E. necator* is able to grow on the surface of all green tissues of the cultivated grapevine *Vitis vinifera* ssp. *vinifera* (*V. vinifera*). The highest damage is caused by infection of unripe berries. At this stage, PM infestation provokes the growing berries to crack open providing entry points for any secondary bacterial and / or fungal infections eventually leading to rotting of the bunches [3,4].

Roughly 170 years ago, *E. necator* was one of the three grapevine-pests introduced to Europe by trading of grapevines derived from crosses of native North American *Vitis* species with *V. vinifera by* England, France and Spain and America [1]. This was the first encounter of *V. vinifera* with this already highly adapted grapevine pathogen on the Eurasian continent explaining the high susceptibility of the cultivated grapevine towards PM. The combination of the pathogenic insect phylloxera (*Daktulosphaïra vitifoliae*), an obligate biotrophic oomycete causing downy mildew (*Plasmopara viticola*; DM) and PM was responsible for the collapse of wine production in France and Spain roughly 150 years ago [1]. The soil borne stage of phylloxera infests the roots of grapevines causing damage and entry points for secondary infections. This results in low yield and eventually in dieback of infested grapevines after several seasons [5]. In addition, the two mildews infect all green tissues of the grapevine. Infections early in the season can lead to complete loss of harvest if DM and PM infect young flowers. The phylloxera-problem was solved by the invention of “crafting” the high wine-quality scions on phylloxera-resistant *Vitis* hybrid rootstocks. Protection against the two mildews was achieved by the invention of the “Bordeaux mixture”, a mixture of Sulphur- and Copper-compounds that prohibits the development of DM and PM when applied prior to infections [6]. This mixture was so effective that even today, 170 years later, it still plays a central role in the plant protection regime of most viticulturists, including organic wine growers. However, to achieve effective plant protection for the highly PM and DM susceptible *V. vinifera* cultivars, fungicides such as the Sulphur- and Copper-compounds or other synthetic protectants have to be applied depending on the environmental conditions up to 12 times during the growing season [7]. This makes viticulture one of the highest agricultural consumers of fungicides [8]. Furthermore, these applications make viticulture laborious and are harmful for humans and the environment due to residues on grape clusters and rain wash-off from plants after treatment [9,10]. On top, an unambiguous correlation of wine growing regions and copper accumulation in top soils was shown. This Copper can be washed off into the nearby rivers and damage non-target organisms [11].

One way to reduce the enormous amounts of fungicides used in viticulture is to breed novel resistant grapevine cultivars carrying resistance traits against DM and PM combined with high wine quality [1,12]. Due to co-evolution of DM and PM with wild *Vitis* species in North America, some accessions of these species have evolved natural genetic resistances which either inhibit the growth of the pathogen partially or completely. In the last decades, roughly 13 of such natural genetic resistance loci against PM have been identified [13–15]. They were delimited to certain regions on various chromosomes of the grapevine genome. Such loci are exploitable by grapevine breeders for introgression into new cultivars with the assistance of MAS. However, it is crucial for breeders to know which resistances to stack to achieve the most durable effect against PM. Therefore, a detailed characterization of the individual resistance loci and their function is essential. This requires artificial inoculation experiments followed by evaluation at different time points of pathogenesis [16].

The resistance locus *Ren9* was identified during a fine-mapping study of the resistance locus *Ren3* on chromosome 15 of ‘Regent’ [15]. It is located in the anterior part of chromosome 15 spanning an interval of roughly 2.4 Mb. To confirm this locus and possibly further delimit the resistance-mediating region on chromosome 15, a cross of ‘Regent’ and ‘Cabernet Sauvignon’ was phenotypically characterized repeatedly throughout the growing season of 2016. In addition, controlled experimental inoculations were performed with selected F_1_ genotypes from that cross that carry meiotic recombinations within chromosome 15. In the frame of this work new genetic insertion / deletion (Indel) markers were designed spanning the previously delimited region for *Ren9* with a spacing of 0.1 – 0.2 Mb. These markers allow a possible further delimitation of the resistance locus *Ren9* on chromosome 15 in the grapevine genome.

## 2. Results

### 2.1 Phenotypic field-data

Phenotypic data from the cross population of ‘Regent’ x ‘Cabernet Sauvignon’ were recorded four times during the growing season 2016. This approach was chosen since previous phenotypic evaluations that had been performed at the end of each season yielded scores of around 5 to 9 for nearly all genotypes (whether they were resistant or susceptible) and were blurring genetic differences due to the late evaluation date. The same approach was applied earlier in the cross population of ‘Regent’ x ‘Lemberger’, which allowed the observation of shifting QTLs during the season [15]. According to their genotypic profiles the F_1_ individuals were grouped in either resistant (*Ren3-Ren9*) or susceptible and individuals with either *Ren3 (“Ren3-* only*”)* or *Ren9 (“Ren9*-only*”)*. The distribution of phenotypic data is visualized in Figure 1. The significance of the difference between resistant und susceptible genotypes is indicated above the boxplots (Figure 1). Differences between *Ren3* and *Ren9* carrying F_1_ individuals were not further investigated due to the fact that these two groups are represented by only two individuals each (Figure 1).

**Figure 1:**
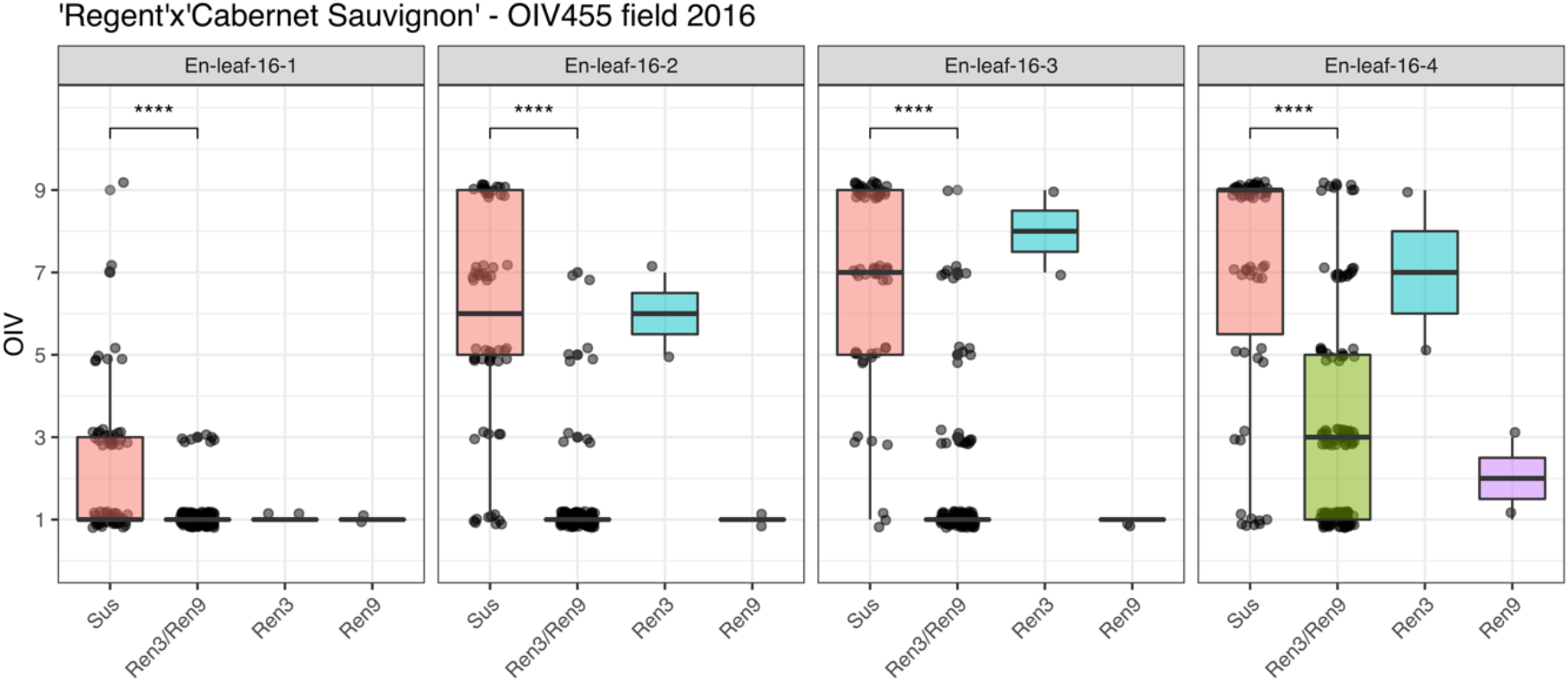
Boxplots of assigned phenotypic scores for the genotypic groups of ‘Regent’ x ‘Cabernet Sauvignon’. Boxes indicate the interquartile range. The median for the respective dataset is indicated by a horizontal line in the boxplot. Number of individuals: susceptible (sus) n=62, Ren3/Ren9 n=132, Ren9 n=2, Ren3 n=2. (*** = P ≤ 0.001, ** = P ≤ 0.01, * = P ≤ 0.05, NS = not significant P > 0.05)

The phenotypic scores in the first scoring date are shifted towards 1 as the medians indicate in the boxplots (Figure 1). The main distribution of phenotypic scores ranged from 1 to 5 in this dataset which was due to the early date of scoring. However, significant differences could be detected between susceptible and resistant genotypes (Figure 1, 16-1: sus – Ren3-Ren9 ***). The median of susceptible genotypes is continuously shifted towards 9 in the three following datasets (Figure 1, 16-2, 16-3, 16-4). For genotypes with *Ren3* and *Ren9* associated alleles the median shifts to 3 in the last dataset which represents the scoring date at the end of the season with highest infection pressure (Figure 1, 16-4). The two individuals with only *Ren3* also show a continuous shift towards score 7, indicating a rather strong infestation with *E. necator* (Figure 1, 16-4). In contrast, for the two individuals carrying *Ren9*, the median score is shifted to score 2 at the last date (Figure 1, 16-4).

### 2.2 QTL-analysis with phenotypic field-data

The described phenotypic data was used for QTL-analysis with the previously published genetic map of ‘Regent’ x ‘Cabernet Sauvignon’ [15]. QTL analysis was performed with the maternal (‘Regent’) and paternal (‘Cabernet Sauvignon’) genetic map. Therefore, the genotypic data was coded as doubled haploid (DH) according to the manual of JoinMap®4.1. Results for the ‘Regent’ haplophase are listed in Table 1 and are shown as graph in Figure 2. The results for the ‘Cabernet Sauvignon’ haplophase are shown in Figure S1. In this haplophase no LOD score higher than 3 was detected and therefore this haplophase was not further investigated. For all scoring dates, a QTL for resistance to powdery mildew was observable on chromosome 15 (Table 1, Figure 2).

**Table 1:** QTL-analysis results for PM resistance scored at four different times of the epidemic (E.n.— leaf-16-1 to 16-4) together with the genetic map of LG15 of ‘Regent’.

|  | Data | Mapping | LOD<sub>max</sub> | % Expl | Nearest Marker | QTL-interval (LOD<sub>max</sub> ±1) | LG15 LOD<br>p≤0,05 | Interval<br>[Mb] |
| --- | --- | --- | --- | --- | --- | --- | --- | --- |
| ‘Regent’<br>LG15 | *E.n.-leaf-16-1* | IM | 11,96 | 23,1 | GF15-30/UDV116 | GF15-62 - GF15-44 | 1,3 | 8,7 |
|  |  | MQM | 11,96 | 23,1 | UDV116 | UDV116 - ScORA7 |  | 3.1 |
|  | *E.n.-leaf-16-2* | IM | 39,47 | 58,6 | CenGen6/CenGen7 | CenGen6 - GF15-10 | 1,2 | 0.8 |
|  |  | MQM | 39,47 | 58,6 | CenGen6 | GF15-62 - CenGen7/6 |  | 0.2/0.5 |
|  | *E.n.-leaf-16-3* | IM | 44,96 | 63,8 | CenGen6/CenGen7 | GF15-62 - CenGen7/6 | 1,3 | 0.2/0.5 |
|  |  | MQM | 44,96 | 63,8 | CenGen6 | GF15-62 - CenGen7/6 |  | 0.2/0.5 |
|  | *E.n.-leaf-16-4* | IM | 19,25 | 34,7 | GF15-62 | GF15-59/58 - GF15-54/55 | 1,2 | 2.7 |
|  |  | MQM | 19,25 | 34,7 | GF15-62 | GF15-59/58 - GF15-62 |  | 0.9 |

**Figure 2:**
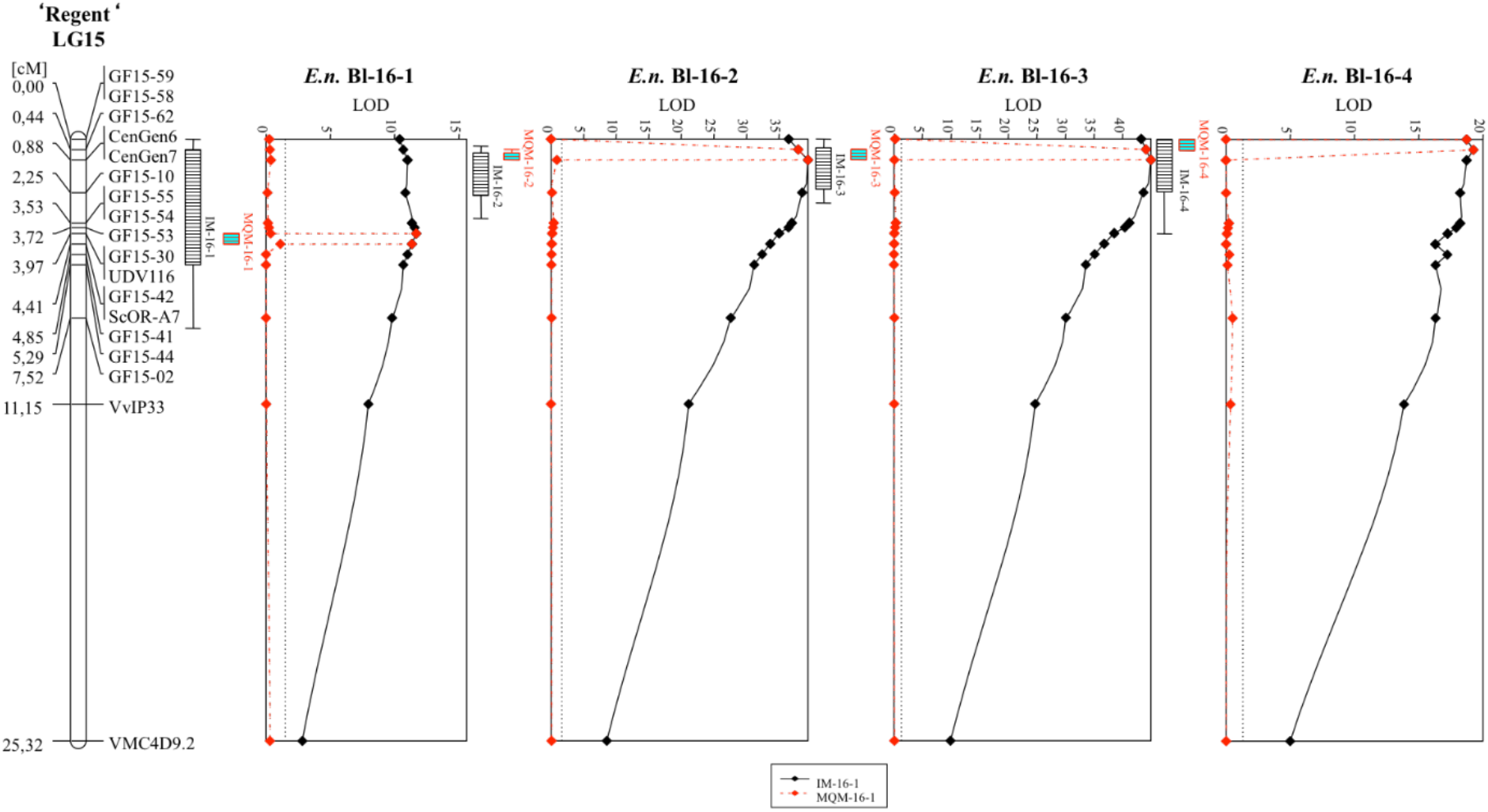
QTL graphs for the four analyzed scoring dates in 2016 with the genetic map of ‘Regent’ derived from the ‘Regent’ x ‘Cabernet Sauvignon’ mapping population. The continuous black line shows the results of IM while the dotted red line indicates the MQM results. The confidence intervals of +/−1 and +/−2 LOD values are indicated by the box and its whiskers at the left side of each graph.

The first scoring (*E.n.*-leaf-16-1) yielded a rather low LOD value of approximately 12 compared to the later three scoring dates (Table 1, Figure 2). This QTL explained around 23% of the observed phenotypic variation. The interval mapping (IM) analysis pointed to an interval spanning the region between markers GF15-62 and GF15-44 (Table 1). This represents around 8.7 Mb of chromosome 15 according to the reference genome PN40024 12X v2. The following MQM mapping limited the region to the interval around UDV116 to ScORA7 (3.1 Mb) with UDV116 being the nearest correlating marker (Table 1, Figure 2). The subsequent scoring dates yield QTLs with LOD_max_ scores of 39 (*E.n.*-leaf-16-2) and 45 (*E.n.*-leaf-16-3) and explained up to 63% of observed phenotypic variation (Table 1). The intervals of the IM analysis were limited to CenGen6 – GF15-10 for *E.n.*-leaf-16-2 (0.8 Mb) and to GF15-62 – CenGen7/6 for *E.n.*-leaf-16-3 (0.2/0.5 Mb). Downstream MQM analysis limited the interval for both scoring dates to the region between GF15-62 and CenGen7/6 representing 0.2 resp. 0.5 Mb on chromosome 15 (Table 1, Figure 2). The forth scoring yielded a QTL, which was shifted completely to the beginning of chromosome 15 (Figure 2). This QTL was represented by a LOD_max_ score of 19 and represented 35% of observed phenotypic variance (Table 1). The interval of this QTL spanned the genetic markers GF15-59/58 and GF15-54/55 that corresponds to 2.7 Mb. Subsequent MQM mapping limited the interval to GF15-59/58 – GF15-62 (Table 1). Taken together, a shift of the QTL from the middle part (*Ren3*) to the anterior part (*Ren9*) of chromosome 15 is observed during the time of beginning of the season to its end.

### 2.3 Fine mapping of the Ren9 region in leaf disc assays

Controlled infection assays were done with leaf discs from selected F_1_ individuals (Table 2) chosen according to their meiotic recombination points on chromosome 15.

**Table 2:**
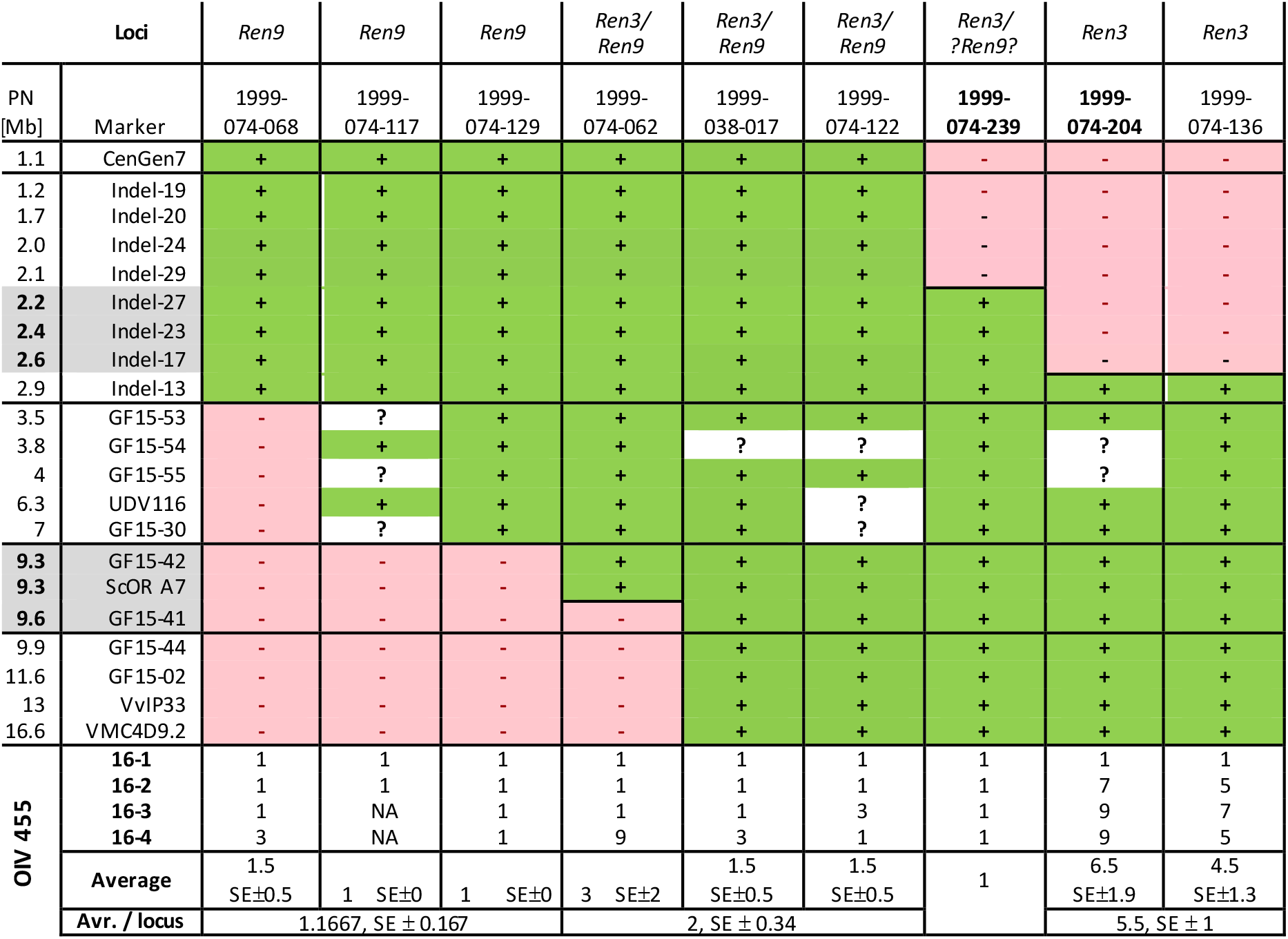
Individuals from the cross ‘Regent’ x ‘Cabernet Sauvignon’ with meiotic recombinations on chromosome 15. SSR-Markers and newly designed Indel-markers are shown: resistance associated allele (+), no resistance associated allele (-), marker not called (?). Together with the recombination points the inverse OIV455 scorings (1 - highly resistant, 9 – highly susceptible) are shown. Genetic markers in regions of *Ren3* ([15], GF15-42, ScOR-A7, GF15-41) and *Ren9* (Indel-27, Indel-23, Indel-17) are marked in grey.

For delimiting the region around *Ren9,* new genetic markers were designed based on insertions and deletions. Table 2 presents the recombination points of the selected F_1_ genotypes from the ‘Regent’ x ‘Cabernet Sauvignon’ cross. Oligonucleotide sequences and amplicons are shown in Sup. Table 1. Individuals with *Ren3*/*Ren9* and “*Ren9-*only” show an average OIV 455 score of 1.7 and 1.16, respectively (Table 2). Assuming the location of the resistance conferring gene of *Ren9* in the interval from CenGen7 to Indel-13, two of the recombinants show only *Ren3* associated alleles. These exhibit an average OIV 455 score of 5.5 (Table 2). In contrast to the two “*Ren3* only” individuals, the F_1_ plant 1999-074-239 shows resistance associated alleles for the markers Indel-27, Indel-23 and Indel-17 and an average phenotypic score of 1.0 (Table 2).

### 2.4 Characterizing the PM single spore isolate GF.En-01

For controlled infection phenotyping, leaf disc inoculation experiments were performed with the aforementioned F_1_ individuals and a single spore PM isolate, GF.En-01. The latter was sampled from a susceptible grapevine cultivar around the JKI Institute for Grapevine Breeding Geilweilerhof, Germany. Genotyping of this isolate showed that it is most likely of EU-B type according to the identified and translated allele sizes described [17] (Table 3). There was some uncertainty for the allele sizes of EnMS-03 and −06 as they differed more than 2 bp from the published sizes (Table 3).

**Table 3:** Allele sizes of the PM isolate GF.En-01 for EnMS markers [27]. Genetic markers with uncertain allele results are marked with a black box. If no allele of corresponding size was found in the list of Frenkel et al., 2012 a ‘?’ was inserted.

|  | EmMS-01 | EmMS-02 | EmMS-03 | EmMS-04 | EmMS-05 | EmMS-06 | EmMS-07 | EmMS-08 | EmMS-09 | EmMS-10 | EmMS-11 |
| --- | --- | --- | --- | --- | --- | --- | --- | --- | --- | --- | --- |
| Allele | 219 | 168 | 218 | 290 | 167 | 249 | 177 | 185 | 155 | 251 | 171 |
| EU-Isolate | A+B | B | B | B? | A+B | B? | B | B | A+B | A+B | B |
| Frenkel et al, 2012 | 239 | 185 | 236 | 305/? | 186 | 266/? | 195 | 205 | 176 | 271 | 191 |
| Frenkel et al, M13 adj. | 220 | 166 | 217 | 286/? | 167 | 247/? | 176 | 186 | 157 | 252 | 172 |

To test the aggressiveness of this isolate, inoculations with *in vitro* plants of ‘Regent’ and ‘Chardonnay’ were performed. Samples were taken one, four, five and 15 days past inoculation with day one providing the reference for the latter. The increase of fungal biomass could be observed for both genotypes. At four dpi a significant difference between ‘Regent’ and ‘Chardonnay’ was observable which was absent at 5 dpi. After 15 days a clear difference between ‘Regent’ and ‘Chardonnay’ was observed with ‘Chardonnay’ showing a median fold change of approximately 65 compared to a fold change of around 20 for ‘Regent’ (Figure 3, A).

**Figure 3:**
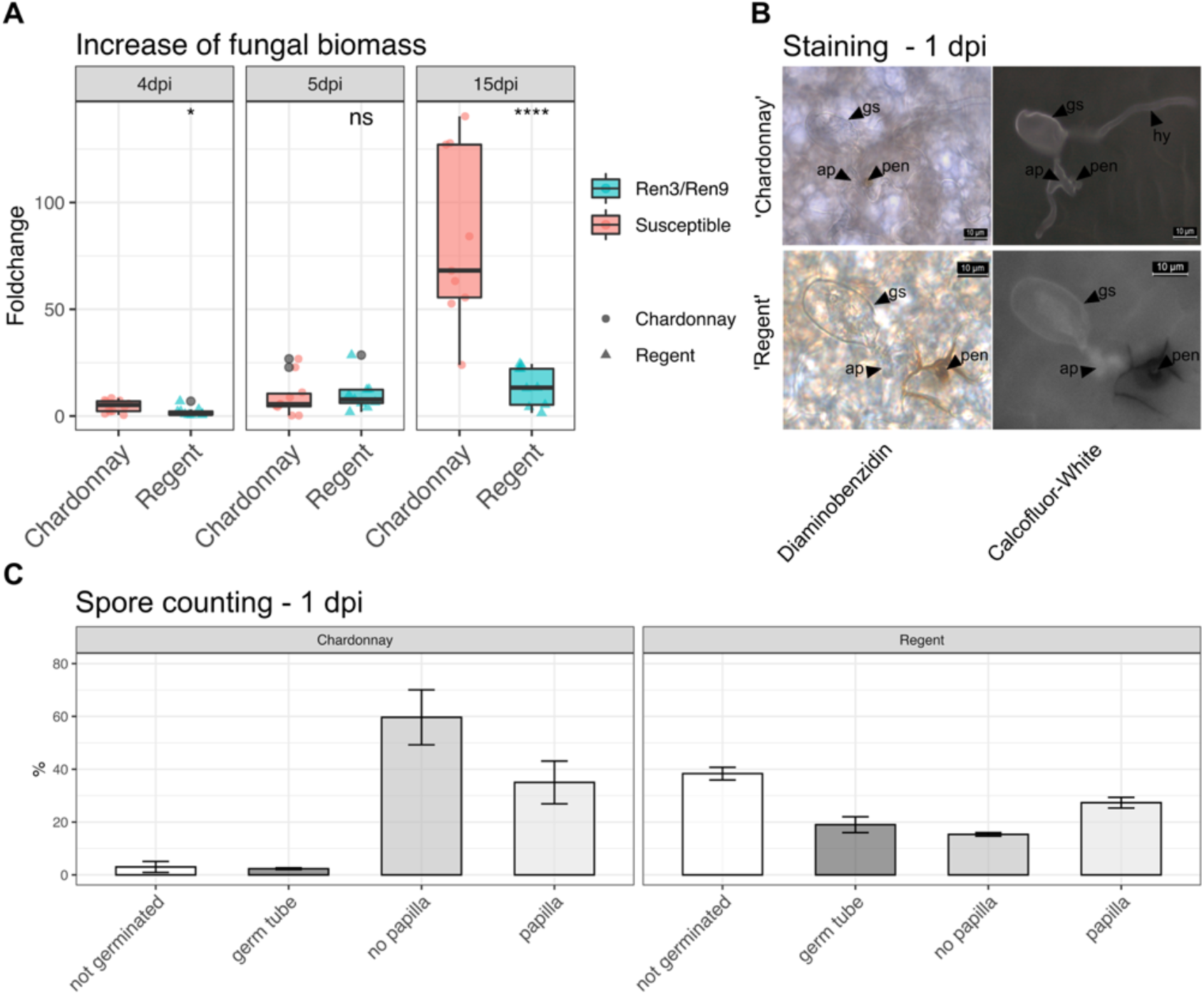
Characterization of PM isolate GF.En-01. **A** Fungal biomass increase over time as measured by qPCR for ‘Chardonnay’ and ‘Regent’. **B** Staining of leaves one day past inoculation with Diaminobenzidin and Calcoflour-White (gs = germinated spore, hy = hyphae, ap = appressoria, pen = penetration site). **C** Counting of conidiospores one day past inoculation and grouping them according to different developmental stages.

In addition, at one day after inoculation, the leaf discs were stained with Diaminobenzidin (DAB) and Calcoflour-White (CW). The DAB stain visualizes reactive oxygen species (ROS) by forming a brown stain at sites with elevated ROS levels. The CW stain visualizes the transparent conidospores and hyphae. A clear accumulation of ROS was observable at the penetration site of the appressoria in ‘Regent’. The brown DAB stain extended around the cell in the appoplast. This reaction was much less pronounced and restricted to the actual penetration site in the susceptible ‘Chardonnay’. Furthermore, primary and secondary hyphae were observed on susceptible ‘Chardonnay’ leaves (Figure 3, B).

During staining spores were counted and grouped according to different developmental stages. The major difference between the susceptible ‘Chardonnay’ and the resistant ‘Regent’ was the overall germination rate, which was 97 % in ‘Chardonnay’versus 62 % on ‘Regent’. On ‘Regent’ leaves, a big portion of germinated spores showed only germ tubes at one day past inoculation (Figure 4). On ‘Chardonnay’ most of the spores germinated and succesfully formed appressoria. No papilla formation was detectable for the biggest proportion of spores (Figure 3, C, ~60 %). On ‘Regent’ the larger portion of germinated spores were accompanied by papilla formation (Figure 3, C, ~30 %). Taken together, these results indicate that *Ren3/Ren9* is capable of restricting the growth of the GF.En-01 isolate. Studying the two resistances independently should therefore be possible with this isolate. However, it indicates that *Ren3/Ren9* mediates only a partial PM resistance against this *E. necator* EU-B type isolate.

**Figure 4:**
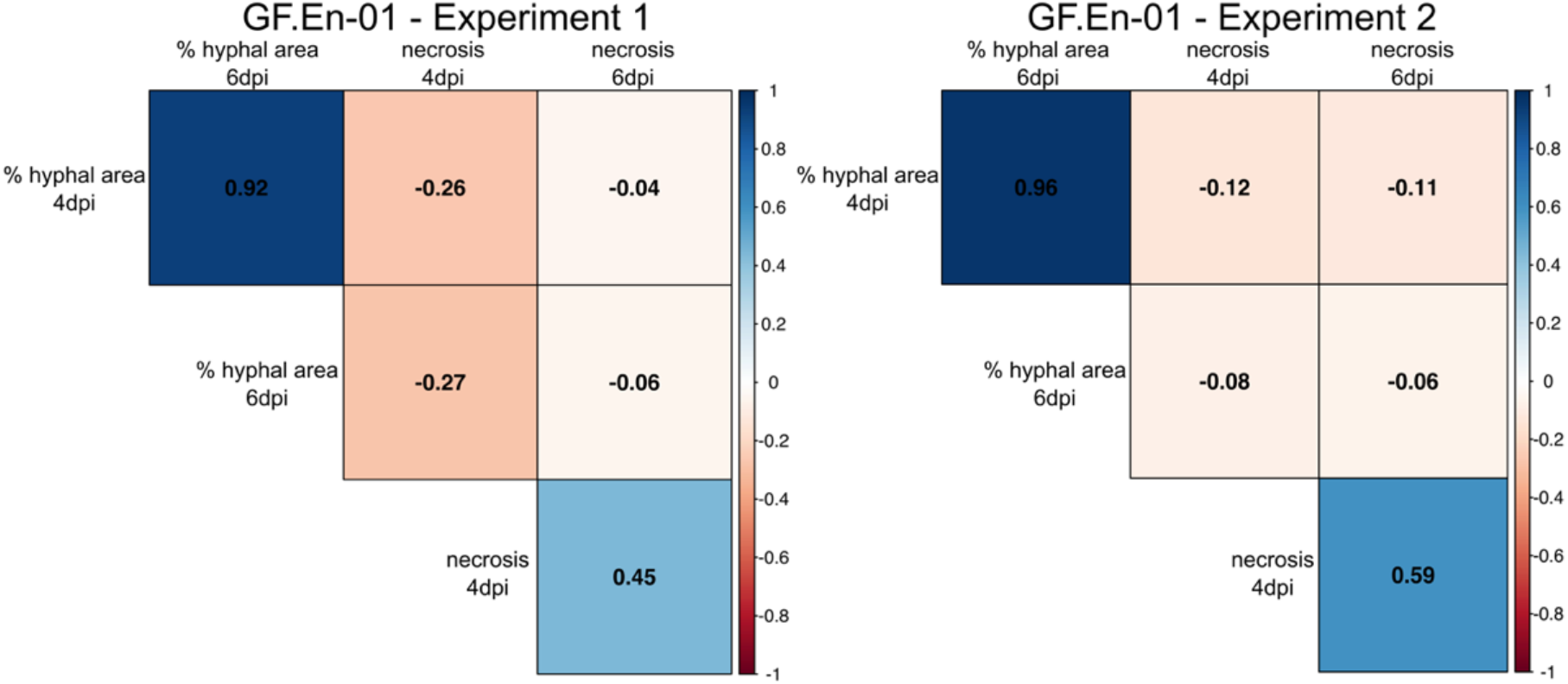
Correlation plot of the percentages of hyphal area and necrosis formation at 4 and 6 dpi. Positive correlations are indicated in blue and negative correlations are indicated with red. The data is split into the two independent experiments. (Significance level: p < 0.05; all correlations were significant)

### 2.5 Leaf disc infection assays with GF.En-01

Two independent inoculation experiments were performed with the single spore PM isolate GF.En-01. Datasets for hyphal growth and necrosis formation from both experiments were compared with each other in a correlation plot. In previous studies a hypersensitive response (HR) / necrosis associated with the appressoria of PM has been proposed as a mechanism for *Ren3* and *Ren9* mediated resistance [15].

Here, a significant positive correlation was observed for percentage of hyphal area present at 4 and 6 dpi comparing both experiments (Figure 4). In addition, a strong positive correlation for necrosis formation was observed for 4 and 6 dpi in both experiments (Figure 4). Percentage of hyphal area showed in all cases a negative correlation with necrosis formation. The strongest negative correlation was observed in both experiments at 4 dpi (Figure 4), indicating a small negative effect of necrosis formation on hyphal growth. Six days past inoculation only a very weak negative correlation was found between these two scored traits (Figure 4).

After a global analysis of the datasets an analysis of the different R-loci combinations was performed by grouping the phenotypic scores of F_1_ individuals from the ‘Regent’ x ‘Cabernet Sauvignon’ cross with similar combinations. As controls, a breeding line with the strong PM resistance locus Run1 and the PM susceptible genotypes ‘Cabernet Sauvignon’, ‘Chardonnay’ (experiment 1, GFEn01_1) and ‘Diana’ (experiment 2, GFEn01_2) were added in the experiments (Figure 5, Sus, Run1). Means of the different R-loci combinations were compared to the susceptible group to detect statistical differences. For all groups a significant difference to the susceptible control could be observed at four- and six-days past inoculation (Figure 5). For Run1, a strong HR was observed associated with the primary appressoria of the conidospores of GF.En-01, as already well documented in several studies [18–20](Figure S2). This HR prevented any growth of PM on leaf discs of this genotype (Figure 5, Run1). In contrast to that, individuals with the different Ren3 and Ren9 combinations showed variable resistance to PM. Phenotypic scores of Ren3/Ren9 individuals showed the highest variation and were overlapping four- and six-days past inoculation with those of the susceptible control group (Figure 5).

**Figure 5:**
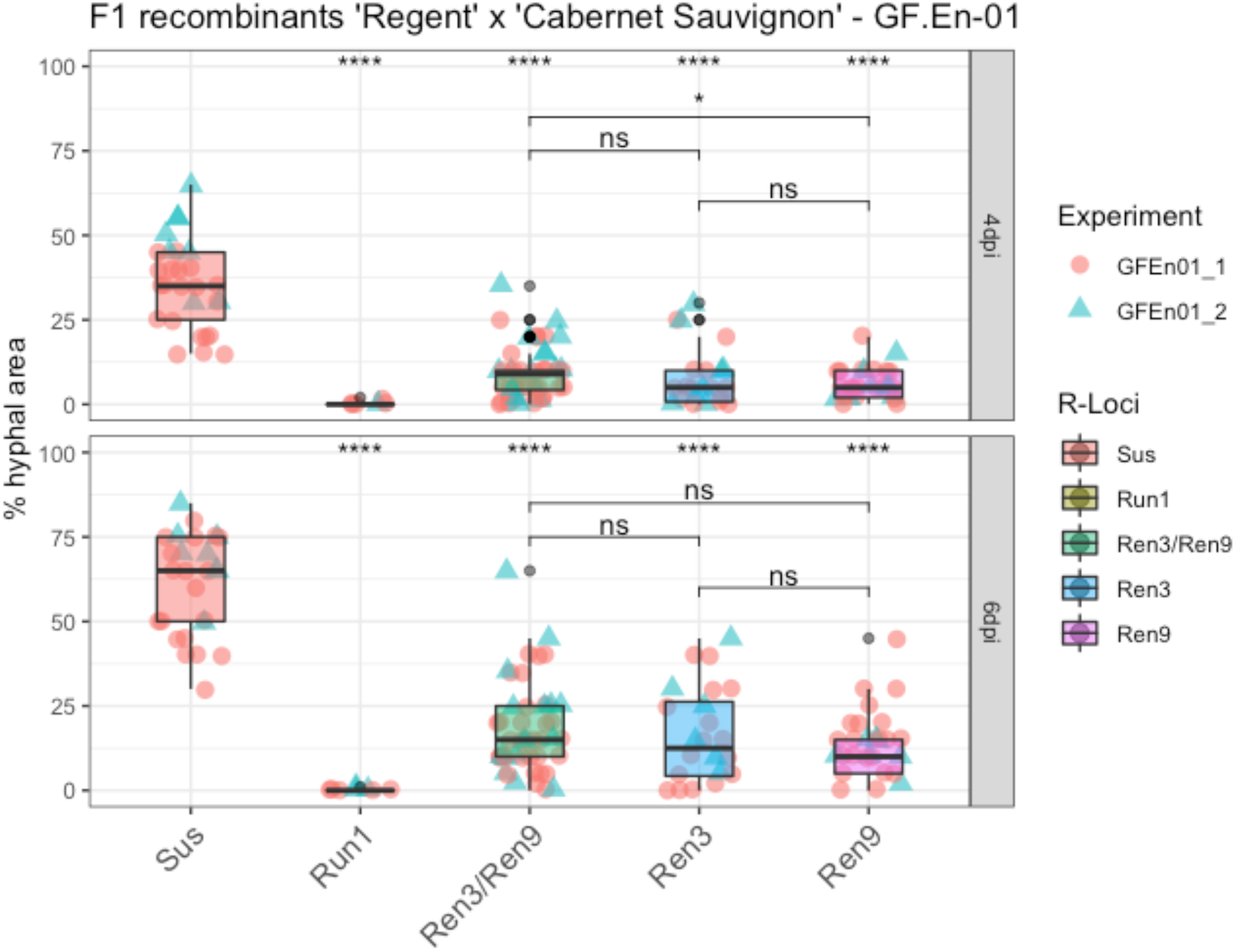
Boxplots of percentage of hyphal area at four (4dpi) and six days past inoculation (6dpi). F_1_ individuals with the same R-locus combination were grouped. Phenotypic scores from different experiments are indicated by different shapes and colors of the data points. Outliers were colored in black. Mean of respective groups are compared to the susceptible group and the mean of the different *Ren3* and *Ren9* combinations with each other (**** = P ≤ 0.0001, *** = P ≤ 0.001, ** = P ≤ 0.01, * = P ≤ 0.05, NS = not significant P > 0.05).

However, the median of percentage hyphal area of the *Ren3/Ren9* group increases from approximately 12 % to roughly 18 %, which is a clear difference compared to the ~35 % to ~65 % change of the susceptible group (Figure 8). To test if there is any significant difference between “*Ren3-only“* or “*Ren9-*only” and the combination of both resistance loci, the means of these groups were compared. Only at 4 dpi a significant lower percentage of hyphal area was observed for “*Ren9-*only” compared to “*Ren3/Ren9”* (Figure 5). After six days, no significant differences were observed between the three groups (Figure 5).

In addition to percentage hyphal area, the trait necrosis formation was scored. The phenotypic data was analyzed the same way as percentage of hyphal area. Necrosis formation of the different *R*-loci combination carriers was compared to the susceptible control group. At both four- and six-days past inoculation the grapevines with the various R-loci combinations showed a significant difference compared to the susceptible group (Figure 6).

**Figure 6:**
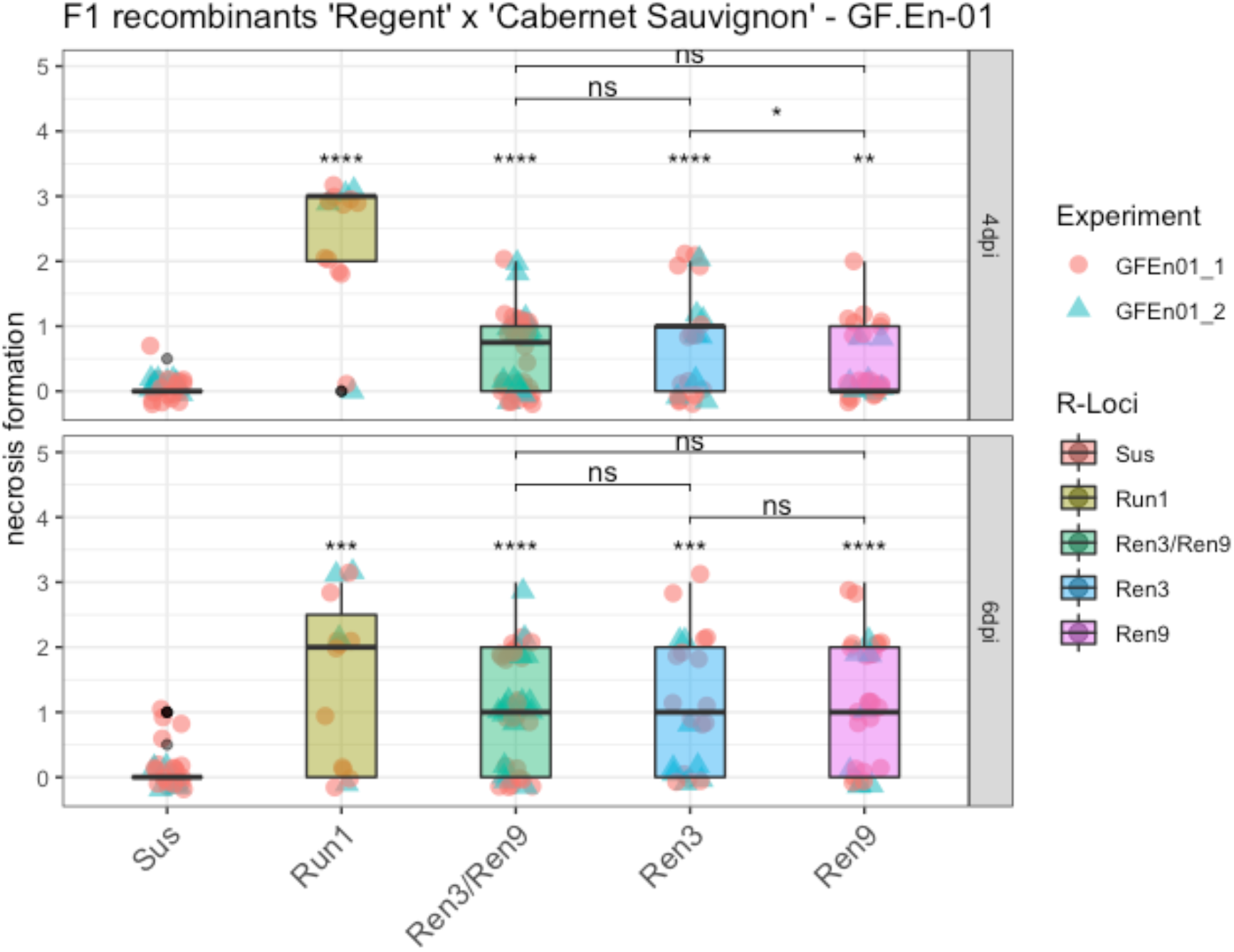
Boxplots of necrosis formation associated with appressoria at four (4dpi) and six days past inoculation (6dpi). F_1_ individuals with the same R-locus combinations were grouped. Phenotypic scores from different experiments are indicated by different shapes and colors of the data points. Outliers were colored in black. Mean of respective groups are compared to the susceptible group and the mean of the different Ren3 and Ren9 combinations with each other (**** = P ≤ 0.0001, *** = P ≤ 0.001, ** = P ≤ 0.01, * = P ≤ 0.05, NS = not significant P > 0.05).

The breeding line with *Run1* showed, as already described, a strong HR associated with nearly all primary appressoria formed by the conidiospores, which is indicated by a median score of three and two at 4 dpi and 6 dpi (Figure 6, *Run1*). Median scores of *Ren3/Ren9* and “*Ren3*-only” were around one at 4 dpi, whereas *Ren9* showed a median score of zero at 4dpi, a significant difference compared to *Ren3/Ren9* (Figure 6). At 6 dpi the different combinations of *Ren3* and *Ren9* all showed a median score of one but overall the scores were ranging from zero to 3 (Figure 6).

## 3. Discussion

Several studies reported a shift of the QTL for resistance to PM on chromosome 15. Van Heerden et al., 2014 showed a LOD_max_ marker CenGen-6 associated with resistance to PM which is located at 1.4 Mb, and a total interval from CenGen-6 to UDV-116 on chromosome 15 (Figure 7, blue bar). In another study, the same research team showed the *Ren3* QTL associated with marker UDV-116, which is located in the middle of chromosome 15. One could argue that, if the marker density in the anterior part of chromosome 15 would have been increased, the QTL would have been possibly shifted further to the beginning of the chromosome [21]. Teh et al. [22] also investigated the resistance *Ren3* with a SNP based genetic map and phenotypic field data. Their interval for resistance to PM ranged from 0.09 to 2.2 Mb on chromosome 15 (Figure 7, orange bar).

**Figure 7:**
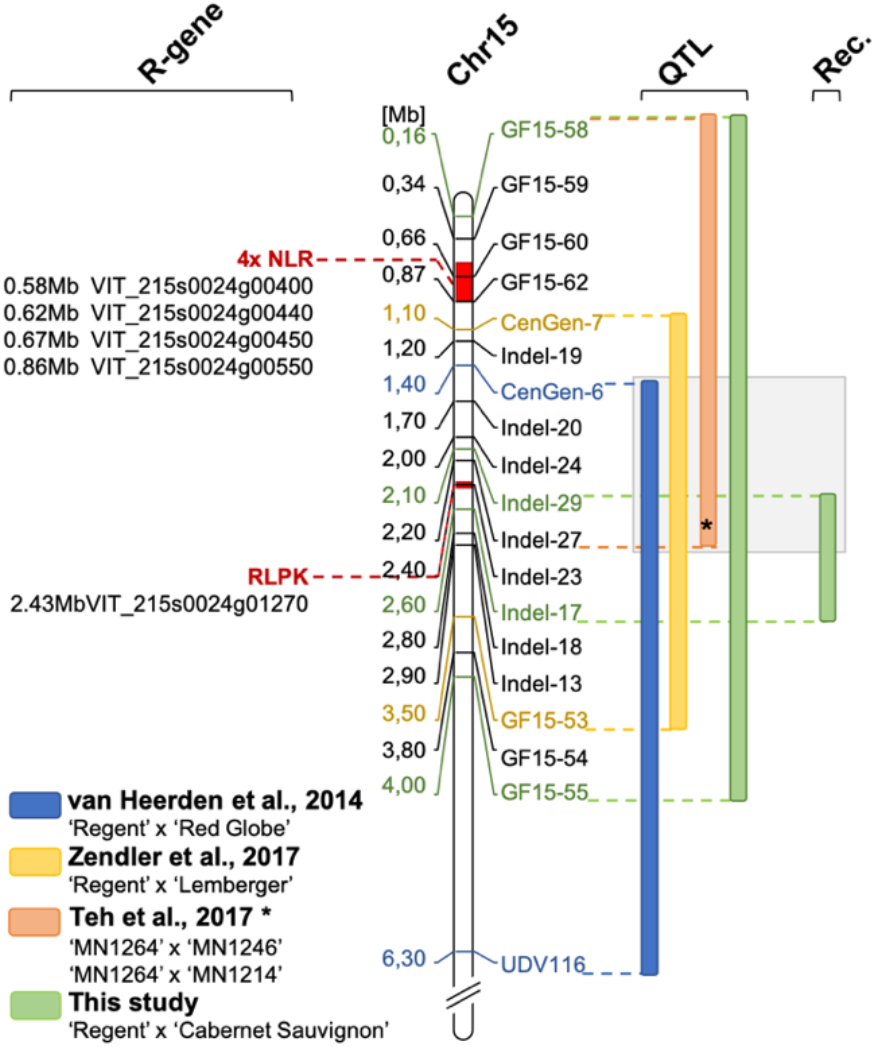
Overview of QTLs for resistance to PM in the anterior part of chromosome 15. Bars next to the map of chromosome 15 indicate QTL intervals (LOD_max_+/− 1). Physical positions are presented in respect to the reference genome of PN40024 12x v2 using the position of the genetic markers applied in this study. The QTL revealed in this study (green bar) represents the largest observed interval in the front part of chromosome 15 (enclosing the results of 16-2, −3 and −4). In addition, the interval resulting from the analysis of F_1_ individuals with meiotic recombination on chromosome 15 is indicated (Rec.). The overlap of all four QTL analyses is highlighted in grey. The region from the start of the chromosome to the position of marker GF15-55 was searched for resistance gene analogs (R-gene, NLR = Nucleotide binding leucin rich repeat, RLPK = receptor like protein kinase). Possible regions are indicated in red. (* The physical position of the QTL interval of Teh et al. [22] was approximated to physical positions of markers in this study according to SNP positions in their supplemental material)

In a previous study for the delimitation of the resistance locus *Ren3*, a second resistance associated region was identified on chromosome 15. This region, now termed *Ren9,* was located in the front part of chromosome 15 and mediated necrosis associated with the appressoria of PM nine days past inoculation [15]. The region of this resistance locus could be delimited to a 2.4 Mb interval flanked by the genetic markers CenGen-7 and GF15-53 (Figure 7, yellow bar) [15].

### 3.1 QTL analysis

In the study presented here a QTL analysis was performed with the previously published genetic map of ‘Regent’ x ‘Cabernet Sauvignon’ [15] and new phenotypic data for resistance to PM. The progression of the infections was scored at four dates during the viticulture season in 2016. Analysis after grouping the individuals into their respective R-loci combinations (susceptible, *Ren3, Ren9, Ren3/Ren9*) indicates a significant difference between the carriers of *Ren3/Ren9* and the susceptible group. At all four scoring dates there is clear evidence for a positive effect of the two R-loci on the inhibition of PM growth (Figure 1). In the beginning of the season, the phenotypes of the majority of the genotypes were shifted towards resistance (Figure 1, 16-1). The QTL results for this date show a QTL-region flanked by the markers UDV-116/GF15-30 and GF15-42/ScOR-A7 (Figure 2). This region agrees with the previously published localization of locus *Ren3* [15,21] and confirms it in this independent mapping study. The phenotypic scores of genotypes without any *R*-locus are shifted towards susceptible (9) from the second scoring date onwards (Figure 1, 16-2 to 16-4) reflecting the developing PM epidemic and increasing infection pressure. The two individuals with only *Ren3* showed a median phenotypic score alike or higher than the susceptible genotypes at the later scoring dates (16-2, 16-3, Figure 1). QTL analysis for the dates 16-2, 16-3 and 16-4 revealed a QTL shift towards the anterior end of chromosome 15 (Figure 2). The flanking markers for the QTLs of dates 16-2 and 16-3 are GF15-62 and CenGen6/CenGen7 (Figure 2, Tab. 1). The LOD_max_ marker in both cases is CenGen6 with a LOD score of 39.5 – 45 explaining 58.6 – 63.8 % of phenotypic variance (Figure 2, Tab. 1). This result agrees with the finding of van Heerden et al. [23], who identified CenGen6 as the left flanking genetic marker in their QTL analysis for PM resistance. QTL analysis with the cross ‘Regent’ x ‘Lemberger’ also had indicated high LOD scores for markers in the anterior part of chromosome 15 for the sampling dates 2015-1, 2015-2 and 2016-1 [15]. The interval mapping of these three dates revealed GF15-10 and CenGen-6 as left flanking genetic markers [15].

The scoring date 16-4 was at the very end of the season and the epidemic. At this time, a strong infection pressure should have been built up resulting in a shift of the phenotypic scores towards susceptibility for all carriers of R-loci (Figure 1). Yet, *Ren3/Ren9* carrying individuals show a median score of 3 and most of the individuals range from 1 – 5 at this time of the season (Figure 2). The QTL analysis with this dataset shows reduced LOD scores for all markers. The QTL region, however, is still associated with the anterior region of chromosome 15. The LOD_max_ marker is GF15-62, indicating a further shift of the QTL region to the beginning of chromosome 15, in agreement with the findings of Teh et al. [22] (Figure 2, Table 1). This marker still explains about 38 % of the observed variance (Table 1). Taken together, these results from four independent grapevine crosses show a high likelihood of *Ren9* being located in the front part of chromosome 15 at around 0 to 4Mb (Figure 7). The region of overlap between all four QTLs ranges from 1.4 to 2.0 Mb defining it as a high confidence area (Figure 7).

Additionally, the new QTL analysis presented here underscores the fact that the loci *Ren3/Ren9* mediate partial-but not total-resistance against powdery mildew.

### 3.2 Fine mapping of the Ren9 region

Detailed investigations were carried out with a subset of individuals exhibiting meiotic recombinations on chromosome 15 that separate the two resistance loci *Ren3* and *Ren9* (Table 2). Newly designed insertion / deletion markers (Indel) are highlighted by a black square around them (Table 2, Table S1). The OIV455 field scores from 2016 are shown and the average was calculated (Table 2). Individuals with the same R-loci combination were grouped and their phenotypic score was averaged. The combined resistance *Ren3/Ren9* or “*Ren9*-only” show an average field score of ~1.2 to ~2 whereas two individuals with “*Ren3-*only” showed an average score of 5.5 (Table 2). One possible explanation for these results might be that during the season in 2016 a change of the composition of PM isolates took place. Isolates that are more virulent may emerge at the end of the season and could be capable of breaking *Ren3*. For Europe two dominant PM isolates have been described termed EU-A and -B [17]. Recent studies on PM isolates in vineyards in Hungary have shown EU-B to be the first isolate in the season sampled on flag-shoots. Later during the season in summer and autumn a mixture of EU-B, -B2 and -A was detectable [24]. Similar events may happen in the vineyards around the Institute for Grapevine Breeding Geilweilerhof, Germany and would explain the results for these F_1_ individuals.

However, one individual (1999-074-239) which was previously classified as “*Ren3-*only” showed an OIV455 score of 1 throughout the season (Table 2). For genetic map construction only GF15-53 (3.5 Mb) and CenGen-7 (1.1 Mb) were available as reliable genetic markers to asses recombination points in this genetic area. For fine mapping of the recombination points, new genetic Indel markers were developed in this study (Table 2, Indel, Table S1). These new genetic markers further defined the recombination points for the individuals 1999-074-239, −204 and −136 (Table 2). For the F_1_ individuals 1999-074-136 and 1999-074-204 the recombination point from susceptible to resistant was located between the markers Indel-17 (2.6 Mb) to Indel-13 (2.9 Mb). For the genotype 1999-074-239 the recombination happened between Indel-29 (2.1 Mb) and Indel-27 (2.2 Mb).

As the phenotypic scores of 1999-074-239 are similar to those of *Ren3/Ren9* and “*Ren9*-only” (Table 2, average OIV score of *Ren9 =* 1.2 and *Ren3/Ren9 =* 2) we hypothesize that this individual is carrying both resistances. This would mean that the interval between Indel-29 and Indel-13 (2.1 – 2.9 Mb) could represent the *Ren9* encoding region and delimit this resistance locus to around 0.8 Mb. For breeders this result means a much smaller introgression required to gain resistance and removal of possible genetic drag. The Indel markers designed here can easily be applied for marker-assisted selection in new breeding programs for stacking multiple resistances in novel grapevine cultivars improved in fungal resistance.

The average OIV scores for the different *R-locus* combinations suggests that under field conditions of the year 2016 *Ren9* was the major resistance against PM. Average OIV scores of 1.2 for *“Ren9-*only*”* individuals and 2 for *Ren3/Ren9* plants clearly differ from *“Ren3-*only” carriers with an average score of 5.5 (Table 2). However, in a study published by an Italian research team *Ren9* carrying genotypes exhibited a reduced level of resistance against PM in unsprayed fields in Italy compared to *“Ren3*-only” and *Ren3/Ren9* individuals [25]. This may indicate a different composition of PM isolates in the fields of Germany and Italy with different virulence levels breaking either the resistances encoded by *Ren3* or *Ren9*. However, this hypothesis should be treated with care. In the study presented here, the number of individuals investigated was limited and the observations were only for one year. Further research with more individuals carrying the different *R-*locus combinations over several years is required to elucidate this observation in more detail.

### 3.3 Leaf disc inoculation

The individuals from Table 2 were submitted to artificial inoculation experiments with a single spore isolate sampled in the field of the JKI, Institute for Grapevine Breeding Geilweilerhof, Germany. The PM isolate GF.En-01 was genotyped with the published SSR markers [17]. The allele combinations obtained from the genotyping indicate that this isolate represents most likely the EU-B type (with some uncertainty remaining for the markers EnMS-04 and −06). This divergence can be explained by the limited precision of capillary electrophoresis and the use of fluorescent dyes that might slightly change the apparent size of amplicons. Further, it was necessary to adapt the published sizes of Frenkel et al. [17] by subtracting the 19 bp of the M13 sequencing tag they used from the amplicon sizes obtained in capillary electrophoresis.

Growth of GF.En-01 was significantly reduced on ‘Regent’ compared to the susceptible genotype ‘Chardonnay’ (Figure 3, A). There is clear evidence that the development was much slower on *Ren3/Ren9* compared to the susceptible control (Figure 3, C). The inhibition of growth was most likely due to the establishment of papilla and ROS at sites of penetration (Figure 4, B). These are typical resistance responses against grapevine powdery mildew [13].

After this characterization, GF.En-01 was used for leaf disc inoculation experiments of the recombinant F_1_ individuals from Table 2. Two independent inoculation experiments were performed yielding similar results. The data for the two traits percentage of hyphal area and necrosis formation were tested for correlation to investigate a possible effect of necrosis formation on hyphal growth. In the two independent experiments a weak, yet significant negative correlation between necrosis formation and hyphal growth could be observed at four days past inoculation. This trend was much weaker six days past inoculation (Figure 4). However, these findings indicate that there is indeed an interaction between these traits showing that necrosis formation contributes to some small extent to the inhibition of PM growth.

To investigate the effects of the different *R*-loci combinations in detail, the phenotypic scores of the individuals with similar *R*-loci were grouped. This grouping showed that there is no significant difference between the respective *Ren3* and *Ren9* combinations in terms of percentage of hyphal area covering the leaf discs except at four dpi in the comparison of *Ren3/Ren9* to “*Ren9-*only”. These two showed a significant difference with *Ren9* showing less hyphal growth. Most of the phenotypic scores are overlapping between *Ren3/Ren9* and “*Ren3-*only” making this difference marginal. Nevertheless, for all *R*-loci combinations a significant reduction in hyphal growth compared to the susceptible controls could be observed at both four- and six-days past inoculation (Figure 5). These results indicate that both resistance loci by themselves are capable to detect the EU-B PM isolate and inhibit its growth. It also shows that both resistances are equally strong and no additive effect can be observed when stacking them, at least when dealing with this specific single spore isolate.

The trait necrosis formation was investigated the same way. The *R*-loci combinations of *Ren3* and *Ren9* showed at both dates a significant difference compared to the susceptible controls. A significant difference among the combinations for “*Ren3-*only” compared to “*Ren9*-only” at four days past inoculation was also shown (Figure 7). This might indicate that the mechanism behind the two resistances differs in terms of detection speed. This difference cannot be observed anymore at six days past inoculation. Trailing necrosis, as it was observed for PM on leaves, is described as a part of ontogenetic resistance of grapevine berries of ‘Chardonnay’ [26,27]. However, all artificial inoculation experiments were performed with young and healthy leaves from the shoot tip and trailing necrosis was absent on the susceptible control leaves from ‘Cabernet Sauvignon’, ‘Chardonnay’ and ‘Diana’. These results indicate that the mechanism observed here differs from the one described for ontogenetically resistant grape berries. Therefore, we propose that the resistance of *Ren3* and *Ren9* relies on a faster detection of PM pointing at specific R-gene interactions.

### 3.4 Possible candidate genes

The resistances *Ren3* and *Ren9* although partial, might rely on different mechanisms. The resistance-associated region for *Ren3* was searched for candidate genes. This yielded a cluster of four NLR genes in the reference genome [15]. Screening the reference genome of PN40024 12x V2 in the proposed QTL interval for *Ren9* yielded two regions with R-gene analogs. The first region, at the very beginning between 0.5 and 0.9 Mb of chromosome 15, comprises a cluster of four possible NLR genes (Figure 7, 4xNLR) and is supported by QTLs from this study and the study of Teh et al. [22]. This cluster might look different in the genome of resistant ‘Regent’. It therefore is of high interest for further investigations. NLR genes have been proven to be key-players in several plant resistance reactions against a multitude of different pathogens [28,29]. Furthermore, the well characterized resistance locus *Run1* which was used in this study as a positive control for resistance against PM was shown to rely on a NLR gene of the “Toll-Interleukin-Receptor-like” type [19].

The second region with another *R*-gene analog is found around 2.4 Mb in a region were multiple QTLs from different crosses overlap ([15,23] and this study). In addition to the QTL intervals, the recombination-points of the F_1_ individuals 1999-074-239 and 1999-074-204 point to this region (Figure 7). The gene found here shows the typical functional domains of a leucin-rich-repeat receptor-like protein-kinase. Such functions are important for the detection of pathogen associated molecular patterns (PAMPs). One of the most prominent examples of PAMP triggered immunity (PTI) is the detection of flagellin by the receptor-like protein-kinase BAK1 in a complex with other receptor like kinases [30]. Roughly 872 of receptor-like kinases are encoded in the grapevine genome [31]. However, any important role of the RLK gene in PM resistance has to be confirmed by functional studies. Therefore, transformations of susceptible cultivars with the possible candidate genes have to be performed and knock-out / -down experiments with resistant cultivars carrying *Ren9* are required. If the RLK gene would prove to be the important one this could indicate that the pathogen perception mediated by *Ren3* and *Ren9* most likely differs between the two. This in turn would be most interesting for breeders. A combination of different resistance mechanisms in pyramiding resistance loci is most promising to generate long-term durability.

## 4. Materials and Methods

### 4.1 Plant material

Progeny used for genetic mapping comprised 236 F_1_ individuals from the cross of ‘Regent’ x ‘Cabernet Sauvignon’. The plants of this population are grown in the experimental fields at JKI Geilweilerhof, Siebeldingen, Germany (49°12’54.1”N 8°02’41.3”E) on their own roots with a spacing of 1.8 to 1.1 m (row by vine). They are cane pruned as is common practice in this wine growing area (Palatinate region). The plantation density at JKI Geilweilerhof, Siebeldingen, is 5050 vines per hectare. The `Regent’ x `Cabernet Sauvignon’ progeny is maintained in an experimental vineyard that was left unsprayed with fungicides. All of the 236 genotypes are represented by one plant each.

For inoculation experiments, plants were kept in the greenhouse as two eye cuttings. Plants were treated with Sulphur once per week to prevent PM infection. One week prior to inoculation experiments plants were not sprayed anymore and the third fully expanded leaf, counting from the shoot tip, was sampled.

*In vitro* plants for artificial inoculation experiments were contained on MS233 (Duchefa, 2.3 g/l) with sucrose (0.11M) and gelrite (0.5 % (w/v), pH 5.8) media. Plants were propagated every 12 weeks by two eye cuttings and were kept in climate chambers with 16 hours light, 8 hours dark and 20 – 22 °C.

### 4.2 Powdery mildew

For controlled inoculation experiments, a single spore isolate was collected from ‘Lemberger’, a susceptible grapevine cultivar grown in the fields of the Institute. The isolate was propagated every three to four weeks on surface sterilized ‘Chardonnay’ leaves maintained on 1% water agar. The inoculated leaves were incubated under long day conditions (16h light, 8h dark). Temperatures were set to 23°C during the day and to 19°C during the night.

### 4.3 DNA extraction

For DNA extraction, about 1cm^2^ pieces of young and healthy leaves were collected from the field and the greenhouse plants, transferred in plastic bags and immediately cooled on ice upon transfer to the laboratory. Leaf segments were shock-frozen in liquid nitrogen and stored at −70°C. DNA was extracted after grinding the samples in the frozen state with a tissue lyser mill (Retsch, 42781 Haan, Germany) using the Macherey Nagel (52355 Düren, Germany) Nucleospin 96 II DNA kit or the PeqGOLD Plant DNA mini Kit (PEQLAB GmbH, 91052 Erlangen, Germany) as described in [14].

### 4.4 Genetic marker design around Ren9

For the development of insertion / deletion (Indel) markers in the *Ren9* region, the reference genome PN40024 12x.v2 [32,33] was used. Genetic marker development was performed as described in Zendler et al. [15]. Sequences showing length polymorphisms greater than six bp were tested for PCR amplification. Unique flanking oligonucleotides for PCR amplification of polymorphic regions were selected according to standard conditions (~50% GC content, 20 – 25 bp lengths, Ta 55 −60°C). PCR reactions were performed in a 10µl reaction mix using the Kapa 2G Multiplex Mix (PeqLAB GmbH, 91052 Erlangen, Germany). PCR products were analyzed on 3% agarose gels with Serva Clear Stain.

### 4.5 SSR-marker analysis

The construction of genetic maps employed SSR markers. SSR marker analysis was performed in multiplex PCR assays with the Kappa2G Multiplex Kit (PeqLAB GmbH, 91052 Erlangen, Germany) mixing up to five different oligonucleotide pairs in one PCR. The forward primer of each pair was 5’-end labeled with fluorescent dyes HEX®, ROX®, FAM® or TAMRA®. Allele sizes were analyzed using the ABI3130XL sequencer with a 36 cm capillary set, a size standard labeled with LIZ® (identical to GeneScan™ 500 LIZ™, Applied Biosystems™) and GeneMapper® 5.0 software (Applied Biosystems™) [15].

### 4.6 Construction of the genetic map

Genetic maps of chromosome 15 were constructed by linkage/recombination analysis using JoinMap®4.1 software [34]. Allele combinations observed in the SSR marker data were encoded according to the manual of JoinMap®4.1. Grouping was done using the independence LOD parameter starting at LOD 2.0 up to LOD 10. Maps were calculated using the Maximum Likelihood mapping algorithm provided in the JoinMap^®^4.1 software. For ‘Regent’ x ‘Cabernet Sauvignon’ the previously published integrated and maternal/paternal genetic maps were used [15]. Individuals which were accidentally selfed or had more than 30% missing data for the analyzed genetic markers were excluded from the mapping calculation to avoid erroneous marker order. Final analysis was based on 236 F_1_ individuals from the cross of ‘Regent’ x ‘Cabernet Sauvignon’.

### 4.7 Phenotypic data

#### 4.7.1 Phenotypic field data

Phenotypic data for QTL analysis was obtained from evaluation of field plants under natural infection pressure with *E. necator*. In former years, resistance scores had been collected once a year in late summer (end of August to end of September) following the inverse OIV (International Organisation of Vine and Wine, http://www.oiv.int) classification as described [14]. In the year 2016, individuals of the cross ‘Regent’ x ‘Cabernet Sauvignon’ were scored four times every three to four weeks (26-06-16 *E.n.*-leaf-16-1, 29-07-16 *E.n.-*leaf-16-2, 18-08-16 *E.n.*-leaf-16-3, 12-10-16 *E.n.*-leaf-16-4). The degree of infection was classified in grades of 1, 3, 5, 7 and 9 (1=no infection at all, 3=nearly no infection visible, 5=punctual infection spots on several leaves, 7=punctual infection on every leaf, 9=infections covering all leaves) (inverse to OIV descriptor 455). Each phenotypic scoring was performed by two people. Scores were assigned by visual inspection of the whole plant.

#### 4.7.2 Artificial inoculation experiments using in vitro plants

For characterization of the *E. necator* single spore isolate GF.En-01, controlled inoculation experiments with *in vitro* plants of ‘Regent’ and ‘Chardonnay’ were performed. Leaves were placed on 1% water agar and inoculated with a brush. Fresh conidiospores were taken from infected ‘Chardonnay’ leaves as described above. To characterize the development of the isolate over time, biomass increase was measured by qPCR [35]. Samples were taken at one-, four-, five- and 15 days past inoculation. A fold change was calculated with the delta-delta ct method, using one dpi as normalization point. Detailed characterization employed Diaminobenzidin (DAB) and Calcofluor-White (CW) staining one day past inoculation. Leaves were first stained by DAB according to a published protocol [36] and then exposed to Calcofluor-White. As the leaves of *in vitro* plants were very tender, the incubation time with DAB was reduced to two hours. For CW staining, leaves were treated according to the manufacturers instructions with one drop of CW staining solution and an equal drop of 10% KOH (Fluka chemicals). The samples were incubated for one minute and then washed with sterile water. Microscopy was performed directly after Calcofluor-White staining. At this point, spores were counted and grouped according to their different developmental stages. For each genotype, three times 100 spores were counted from three independent leaves.

#### 4.7.3 Experimental leaf disc inoculation experiments

For detailed investigations of selected recombinants from the ‘Regent’ x ‘Cabernet Sauvignon’ cross artificial inoculations of leaf discs were carried out as described in [37]. In total, four leaf-discs per genotype from four different plants were placed on 1% water agar plates. They were inoculated with a spore suspension of an *Erysiphe necator* isolate originating from a susceptible ‘Lemberger’ plant in the field.

The PM isolate GF.En-01 was cultivated on leaves of the susceptible cultivar ‘Chardonnay’. Around 10 to 15 days prior to the inoculation experiment six leaves from ‘Chardonnay’ were surface sterilized in 1:10 diluted bleach solution (Eau de Javel, 100ml solution containing 2.6g NaClO) for two minutes. Leaves were rinsed three times with deionized water and dried between paper towels before they were placed in Petri dishes containing 1% water agar. Leaves were then inoculated using 10 to 15 single spore chains of GF.En-01. Leaves for the inoculation experiment were surface sterilized in the same way before punching discs with a one cm diameter cork-borer. The day after placing the leaf discs on 1% water agar the spore suspension was prepared by shaking the inoculated ‘Chardonnay’ leaves in 15 ml sterile water with 10 µl Tween-20. Spores were counted using a hemocytometer. Spore suspensions with 1×10^5^ – 2×10^5^ spores/ml were used for inoculation with a pump sprayer. Visual inspection ascertained that all leaf discs were covered equally with the spore suspension. As a control the cross parental types ‘Regent’ and ‘Cabernet Sauvignon’ were included together with the susceptible cultivar ‘Chardonnay’ or ‘Diana’ as well as a breeding line that carries the strong PM resistance locus *Run1* (VRH3082-1-42).

Leaf disc scoring was performed at four- and six-days past inoculation. The percentage of leaf disc area covered by hyphae and necrosis formation was scored visually using a stereo microscope (Zeiss Axiozoom V16).

Necrosis formation associated with appressoria was scored on a scale of 0 to 3: 0 = no necrosis, 1 = random necrosis associated with appressoria, 2 = trailing necrosis at primary hyphae, 3 = necrosis associated with nearly all appressoria formed by PM (Figure 8). Phenotypic data was analyzed and visualized with R [38] and the packages ggpubr (stat_compare_means()) [39]. Correlations were calculated using the cor() and cor.mtest() package (method = “spearman”) and visualized with the package corrplot() of R [40]. Code and data for reproduction of graphs can be found in the supplemental material (Table S2 – S4).

**Figure 8:**
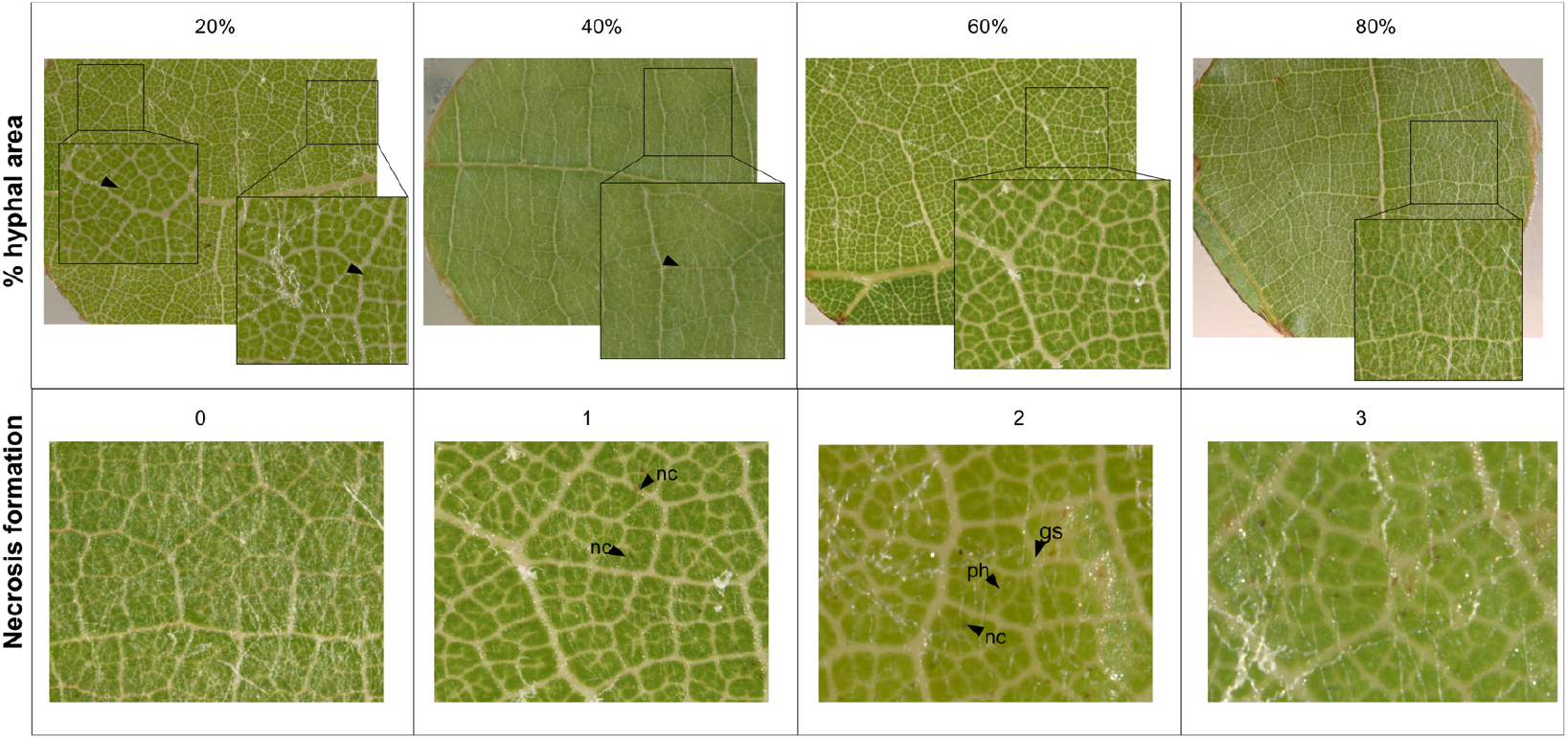
Examples for visual scoring of percentage of hyphal area- and necrosis (gs = germinated spore, ph = primary hyphae, nc = necrosis)

### 4.8 QTL-analysis

QTL analysis was performed using MapQTL®6.0 software [41] with standard settings on the integrated and parental maps. The dataset for the separate parental maps was re-coded as doubled haploid population as recommended in the MapQTL^®^6 manual to avoid “Singularity errors” [41] and enable downstream QTL analysis. The improved maternal genetic map of ‘Regent’ and the paternal genetic map of ‘Cabernet Sauvignon’ were combined with the phenotypic data from the field. Interval mapping (IM) and multiple QTL mapping (MQM) with automatic co-factor selection were performed with the datasets *E.n.*-leaf-16-1, *E.n.-*leaf-16-2, *E.n.*-leaf-16-3 and *E.n.*-leaf-16-4. A permutation test with 1000 permutations determined the linkage group (LG) specific significance threshold for each trait at p ≤ 0.05.

## 5. Conclusions

Both analyses from field and laboratory show that the resistance loci *Ren3* and *Ren9* mediate partial resistance to PM. In generating new breeding lines for European viticulture, *Ren3* and *Ren9* should be complemented by strong resistance loci such as *Run1*, which completely inhibits the progression of the isolate GF.En-01 (representing the EU-B type of powdery mildew). The resistance loci *Ren3* and *Ren9* are broken in Eastern North American vineyards as shown by inoculation experiments with NY19, an PM isolate sampled from vineyards in New York, USA [22]. In controlled inoculation experiments with this isolate Teh and collaborators [22] could not reproduce the QTL from field data for *Ren3/Ren9*. A similar fate might await these resistance loci at some point in Europe, as the evolution of the pathogen never stops. However, the results presented here show that both resistance loci are still useful. Furthermore, the possibility of different mechanisms behind the perception of the pathogen make these resistances very interesting for breeders. With the genetic markers presented here, breeders can easily track the resistance locus *Ren9* in further breeding lines.

## Supplementary Materials

**Figure S 1:**
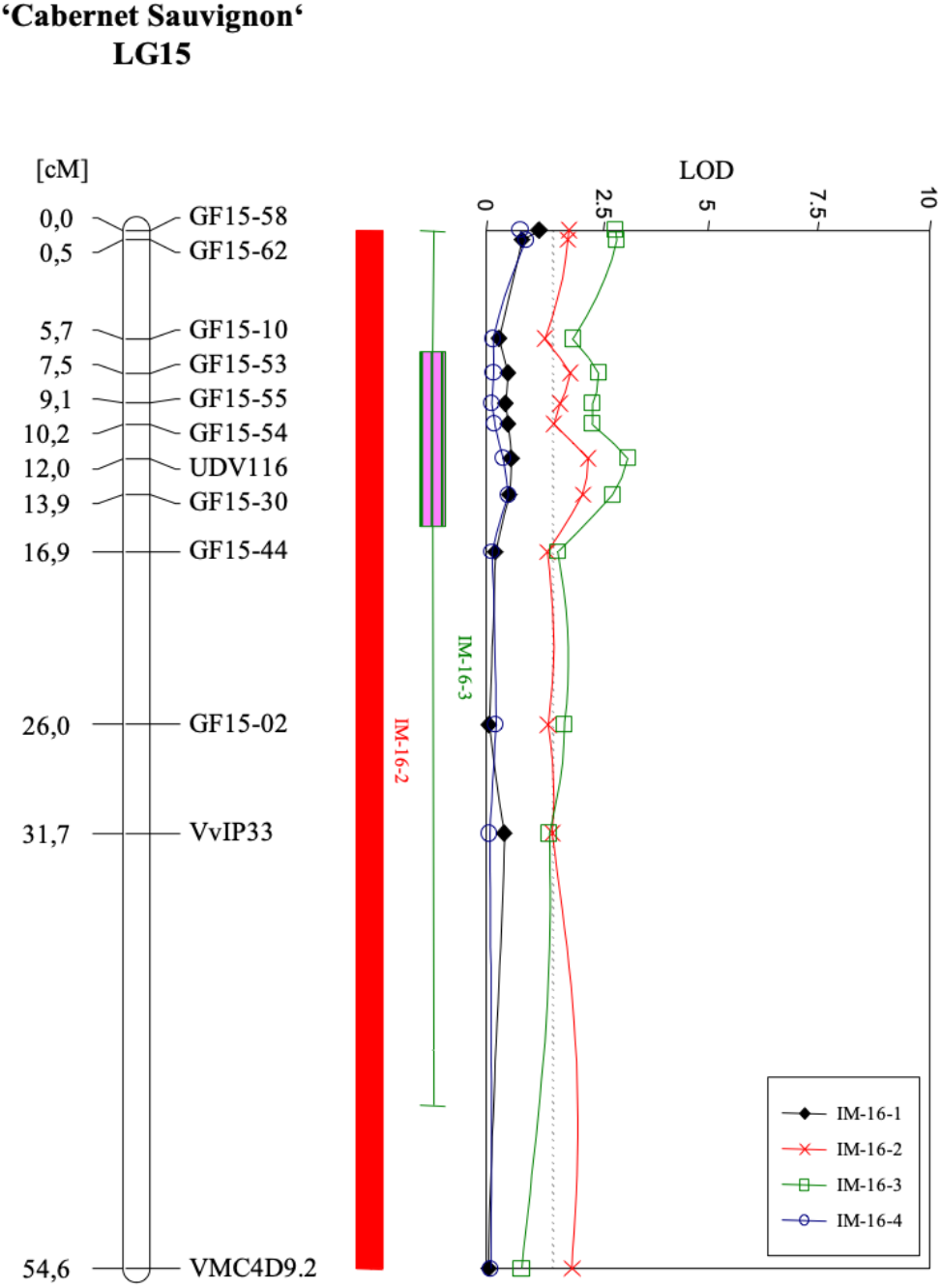
QTL-analysis results for the ‘Cabernet Sauvignon’ haplophase. Results for all four field-scorings are plotted in one graph. The genetic map is given on the left. Significance threshold is 1.2 −1.3 as it was for the ‘Regent’ haplophase. LOD_max_±1 and 2 confidence intervals are indicated by boxes and their whiskers next to the QTL-graph.

**Figure S 3:**
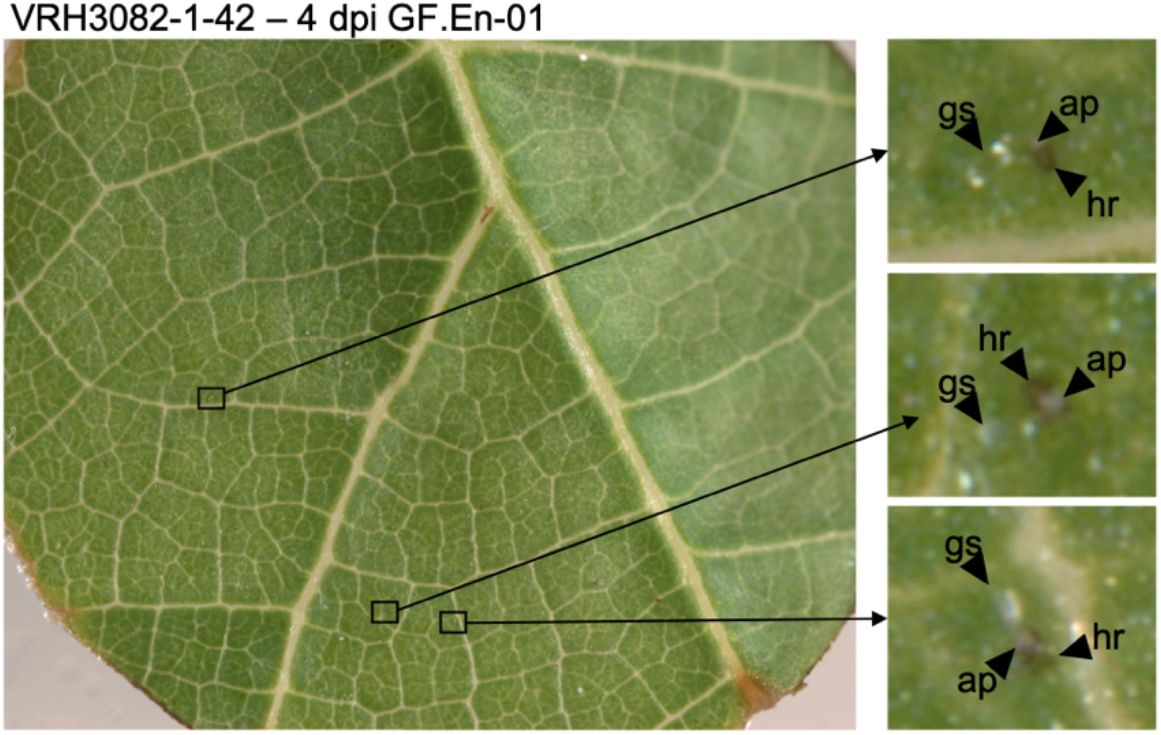
Leaf disc of VRH3082-1-42 (*Run1*) inoculated with GF.En-01 four days past inoculation. Magnified areas show germinated conidiospores (gs) with appressoria (ap) and a hypersensitive response (hr) associated with it.

Table S1: Primer sequences used to amplify the newly designed Indel-maker. Together with the primer sequences amplicon lengths and sequences of Regent (Cham=‘Chambourcin’ haplophase, Dia=‘Diana’ haplophase’, PN40024 12×v2 and ‘Cabernet Sauvignon’ v1.1 are given.

Table S2: Results of artificial inoculations of leaf discs with PM isolate GF.En-01. This table can be used with the supplied R script to reproduce the graph.

Table S3: Data used for correlation plots. This data is for the first experiment (GFEn-01_1). Use the supplied R script to reproduce the graph.

Table S4: Data used for correlation plots. This data is for the first experiment (GFEn-01_2). Use the added R script to reproduce the graph.

Table S5: Data used for spore count plot to characterize GF.En-01 on ‘Regent’ and ‘Chardonnay’.

Table S6: Data used to generate the fungal biomass increase plot to characterize the growth of GF.En-01 on ‘Regent’ and ‘Chardonnay’.

Table S7: OIV455 field scoring data from 2016 used in Figure 2.

## Author Contributions

D.Z. formal analysis, investigation, methodology, visualization and writing (original draft). R.T. resources, writing (review & editing). E.Z. funding acquisition, planning, supervision, writing (review & editing). All authors have read and approved the final manuscript.

## Acknowledgments

Margit Schneider, Sissy Schatt and Claudia Welsch contributed expert technical assistance. We wish to thank Margit Harst and Charlotte Mock for the provision of *in vitro* plants.

## References

1. Töpfer, R.; Hausmann, L.; Harst, M.; Maul, E.; Zyprian, E.; Eibach, R. New Horizons for Grapevine Breeding. In Fruit, Vegetable and Cereal Science and Biotechnology; 2011; pp. 79–96 ISBN 978-4-903313-75-7.

2. Dry, I.; Riaz, S.; Fuchs, M.; Sosnowski, M.; Thomas, M. Scion Breeding for Resistance to Biotic Stresses. In The Grape Genome; Cantu, D., Walker, M.A., Eds.; Springer, 2019; pp. 319–347.

3. Gadoury, D.M.; Cadle-Davidson, L.; Wilcox, W.F.; Dry, I.B.; Seem, R.C.; Milgroom, M.G. Grapevine powdery mildew (*Erysiphe necator*): A fascinating system for the study of the biology, ecology and epidemiology of an obligate biotroph. Mol. Plant Pathol. 2012, 13, 1–16, doi:10.1111/j.1364-3703.2011.00728.x.

4. Wilcox, W.F.; Gubler, W.D.; Uyemoto, J.K. PART I: Diseases Caused by Biotic Factors. In Compendium of Grape Diseases, Disorders, and Pests, Second Edition; Wilcox, W.F., Gubler, W.D., Uyemoto, J.K., Eds.; The American Phytopathological Society, 2015; pp. 17–146.

5. Powell, K.S. A Holistic Approach to Future Management of Grapevine Phylloxera. In Arthropod Management in Vineyards: Pests, Approaches, and Future Directions; Springer Netherlands: Dordrecht, 2012; pp. 219–251 ISBN 9789400740327.

6. Johnson, G.F. The Early History of Copper Fungicides. Agric. Hist. 1935, 9, 67–79.

7. Chen, M.; Brun, F.; Raynal, M.; Makowski, D. Delaying the first grapevine fungicide application reduces exposure on operators by half. Sci. Rep. 2020, 10, 1–12, doi:10.1038/s41598-020-62954-4.

8. Commission, E. The use of plant protection products in the European Union Data 1992-2003 2007 edition; 2007; ISBN 92-79-03890-7.

9. García-Esparza, M.A.; Capri, E.; Pirzadeh, P.; Trevisan, M. Copper content of grape and wine from Italian farms. Food Addit. Contam. 2006, 23, 274–280, doi:10.1080/02652030500429117.

10. Provenzano, M.R.; El Bilali, H.; Simeone, V.; Baser, N.; Mondelli, D.; Cesari, G. Copper contents in grapes and wines from a Mediterranean organic vineyard. Food Chem. 2010, 122, 1338–1343, doi:10.1016/j.foodchem.2010.03.103.

11. Ballabio, C.; Panagos, P.; Lugato, E.; Huang, J.H.; Orgiazzi, A.; Jones, A.; Fernández-Ugalde, O.; Borrelli, P.; Montanarella, L. Copper distribution in European topsoils: An assessment based on LUCAS soil survey. Sci. Total Environ. 2018, 636, 282–298, doi:10.1016/j.scitotenv.2018.04.268.

12. Pertot, I.; Caffi, T.; Rossi, V.; Mugnai, L.; Hoffmann, C.; Grando, M.S.; Gary, C.; Lafond, D.; Duso, C.; Thiery, D.; et al. A critical review of plant protection tools for reducing pesticide use on grapevine and new perspectives for the implementation of IPM in viticulture. Crop Prot. 2017, 97, 70–84, doi:10.1016/j.cropro.2016.11.025.

13. Qiu, W.; Feechan, A.; Dry, I. Current understanding of grapevine defense mechanisms against the biotrophic fungus (*Erysiphe necator*), the causal agent of powdery mildew disease. Hortic. Res. 2015, 2, doi:10.1038/hortres.2015.20.

14. Zyprian, E.; Ochßner, I.; Schwander, F.; Šimon, S.; Hausmann, L.; Bonow-Rex, M.; Moreno-Sanz, P.; Grando, M.S.; Wiedemann-Merdinoglu, S.; Merdinoglu, D.; et al. Quantitative trait loci affecting pathogen resistance and ripening of grapevines. Mol. Genet. Genomics 2016, 291, 1573–1594, doi:10.1007/s00438-016-1200-5.

15. Zendler, D.; Schneider, P.; Töpfer, R.; Zyprian, E. Fine mapping of Ren3 reveals two loci mediating hypersensitive response against Erysiphe necator in grapevine. Euphytica 2017, 213, 68, doi:10.1007/s10681-017-1857-9.

16. Cadle-Davidson, L.; Londo, J.; Martinez, D.; Sapkota, S.; Gutierrez, B. From Phenotyping to Phenomics: Present and Future Approaches in Grape Trait Analysis to Inform Grape Gene Function. In The Grape Genome; 2019; pp. 199–222.

17. Frenkel, O.; Portillo, I.; Brewer, M.T.; Péros, J.P.; Cadle-Davidson, L.; Milgroom, M.G. Development of microsatellite markers from the transcriptome of *Erysiphe necator* for analysing population structure in North America and Europe. Plant Pathol. 2012, 61, 106–119, doi:10.1111/j.1365-3059.2011.02502.x.

18. Agurto, M.; Schlechter, R.O.; Armijo, G.; Solano, E.; Serrano, C.; Contreras, R.A.; Zúñiga, G.E.; Arce-Johnson, P. RUN1 and REN1 Pyramiding in Grapevine (*Vitis vinifera* cv. Crimson Seedless) Displays an Improved Defense Response Leading to Enhanced Resistance to Powdery Mildew (*Erysiphe necator*). Front. Plant Sci. 2017, 8, 1–15, doi:10.3389/fpls.2017.00758.

19. Feechan, A.; Anderson, C.; Torregrosa, L.; Jermakow, A.; Mestre, P.; Wiedemann-Merdinoglu, S.; Merdinoglu, D.; Walker, A.R.; Cadle-Davidson, L.; Reisch, B.; et al. Genetic dissection of a TIR-NB-LRR locus from the wild North American grapevine species *Muscadinia rotundifolia* identifies paralogous genes conferring resistance to major fungal and oomycete pathogens in cultivated grapevine. Plant J. 2013, 76, 661–674, doi:10.1111/tpj.12327.

20. Feechan, A.; Kocsis, M.; Riaz, S.; Zhang, W.; Gadoury, D.M.; Walker, M.A.; Dry, I.B.; Reisch, B.; Cadle-Davidson, L. Strategies for RUN1 Deployment Using RUN2 and REN2 to Manage Grapevine Powdery Mildew Informed by Studies of Race Specificity. Phytopathology 2015, 105, 1104–1113, doi:10.1094/PHYTO-09-14-0244-R.

21. Veikondis, R.; Burger, P.; Vermeulen, A.K.; Van Heerden, C.J.; Prins, R. Confirmation of the effectiveness and genetic positions of disease resistance loci in ‘Kishmish Vatkana’ (*Ren1*) and ‘Villard Blanc’ (*Ren3* and *Rpv3*). South African J. Enol. Vitic. 2018, 39, 185–195, doi:10.21548/39-2-2685.

22. Teh, S.L.; Fresnedo-Ramírez, J.; Clark, M.D.; Gadoury, D.M.; Sun, Q.; Cadle-Davidson, L.; Luby, J.J. Genetic dissection of powdery mildew resistance in interspecific half-sib grapevine families using SNP-based maps. Mol. Breed. 2017, 37, 1, doi:10.1007/s11032-016-0586-4.

23. van Heerden, C.J.; Burger, P.; Vermeulen, A.; Prins, R. Detection of downy and powdery mildew resistance QTL in a ‘Regent’ × ‘RedGlobe’ population. Euphytica 2014, 200, 281–295, doi:10.1007/s10681-014-1167-4.

24. Csikós, A.; Németh, M.Z.; Frenkel, O.; Kiss, L.; Váczy, K.Z. A fresh look at grape powdery mildew (*Erysiphe necator*) a and b genotypes revealed frequent mixed infections and only b genotypes in flag shoot samples. Plants 2020, 9, 1–12, doi:10.3390/plants9091156.

25. Zini, E.; Dolzani, C.; Stefanini, M.; Gratl, V.; Bettinelli, P.; Nicolini, D.; Betta, G.; Dorigatti, C.; Velasco, R.; Letschka, T.; et al. R-Loci Arrangement Versus Downy and Powdery Mildew Resistance Level: A Vitis Hybrid Survey. Int. J. Mol. Sci. 2019, 20, 3526, doi:10.3390/ijms20143526.

26. Gadoury, D.M.; Seem, R.C.; Ficke, A.; Wilcox, W.F. Ontogenic resistance to powdery mildew in grape berries. Phytopathology 2003, 93, 547–555, doi:10.1094/PHYTO.2003.93.5.547.

27. Ficke, A.; Gadoury, D.M.; Seem, R.C.; Godfrey, D.; Dry, I.B. Host barriers and responses to *Uncinula necator* in developing grape berries. Phytopathology 2004, 94, 438–445, doi:10.1094/PHYTO.2004.94.5.438.

28. Lolle, S.; Stevens, D.; Coaker, G. Plant NLR-triggered immunity: from receptor activation to downstream signaling. Curr. Opin. Immunol. 2020, 62, 99–105.

29. Baggs, E.; Dagdas, G.; Krasileva, K. V. NLR diversity, helpers and integrated domains: making sense of the NLR IDentity. Curr. Opin. Plant Biol. 2017, 38, 59–67, doi:10.1016/j.pbi.2017.04.012.

30. Heese, A.; Hann, D.R.; Gimenez-Ibanez, S.; Jones, A.M.E.; He, K.; Li, J.; Schroeder, J.I.; Peck, S.C.; Rathjen, J.P. The receptor-like kinase SERK3/BAK1 is a central regulator of innate immunity in plants. Proc. Natl. Acad. Sci. U. S. A. 2007, 104, 12217–12222, doi:10.1073/pnas.0705306104.

31. Zhu, K.; Wang, X.; Liu, J.; Tang, J.; Cheng, Q.; Chen, J.G.; Cheng, Z.M. The grapevine kinome: Annotation, classification and expression patterns in developmental processes and stress responses. Hortic. Res. 2018, 5, doi:10.1038/s41438-018-0027-0.

32. Jaillon, O.; Aury, J.M.; Noel, B.; Policriti, A.; Clepet, C.; Casagrande, A.; Choisne, N.; Aubourg, S.; Vitulo, N.; Jubin, C.; et al. The grapevine genome sequence suggests ancestral hexaploidization in major angiosperm phyla. Nature 2007, 449, 463–467, doi:10.1038/nature06148.

33. Canaguier, A.; Grimplet, J.; Di Gaspero, G.; Scalabrin, S.; Duchêne, E.; Choisne, N.; Mohellibi, N.; Guichard, C.; Rombauts, S.; Le Clainche, I.; et al. A new version of the grapevine reference genome assembly (12X.v2) and of its annotation (VCost.v3). Genomics Data 2017, 14, 56–62, doi:10.1016/j.gdata.2017.09.002.

34. Van Ooijen, J.W. JoinMap® 4, Software for the calculation of genetic linkage maps in experimental populations 2006.

35. Jones, L.; Riaz, S.; Morales-Cruz, A.; Amrine, K.C.; McGuire, B.; Gubler, W.D.; Walker, M.A.; Cantu, D. Adaptive genomic structural variation in the grape powdery mildew pathogen, *Erysiphe necator*. BMC Genomics 2014, 15, 1081, doi:10.1186/1471-2164-15-1081.

36. Daudi, A. Detection of Hydrogen Peroxide by DAB Staining in *Arabidopsis* Leaves. Bio protoc 2016, 2, 4–7.

37. Cadle-Davidson, L.; Gadoury, D.; Fresnedo-Ramirez, J.; Yang, S.; Barba, P.; Sun, Q.; Demmings, E.M.; Seem, R.C.; Schaub, M.; Nowogrodzki, A.; et al. Lessons from a phenotyping center revealed by the genome-guided mapping of powdery mildew resistance loci. Phytopathology 2016, 106, 1159–1169, doi:10.1094/PHYTO-02-16-0080-FI.

38. R Core Team R: A Language and Environment for Statistical Computing 2019.

39. Kassambara, A. ggpubr: “ggplot2” Based Publication Ready Plots 2020.

40. Wei, T.; Simko, V. R package “corrplot”: Visualization of a Correlation Matrix 2017.

41. Van Ooijen, J.W. MapQTL ® 6, Software for the mapping of quantitative trait loci in experimental populations of diploid species 2009.

